# Disentangling biological and analytical factors that give rise to outlier genes in phylogenomic matrices

**DOI:** 10.1101/2020.04.20.049999

**Authors:** Joseph F. Walker, Xing-Xing Shen, Antonis Rokas, Stephen A. Smith, Edwige Moyroud

## Abstract

The genomic data revolution has enabled biologists to develop innovative ways to infer key episodes in the history of life. Whether genome-scale data will eventually resolve all branches of the Tree of Life remains uncertain. However, through novel means of interrogating data, some explanations for why evolutionary relationships remain recalcitrant are emerging. Here, we provide four biological and analytical factors that explain why certain genes may exhibit “outlier” behavior, namely, rate of molecular evolution, alignment length, misidentified orthology, and errors in modeling. Using empirical and simulated data we show how excluding genes based on their likelihood or inferring processes from the topology they support in a supermatrix can mislead biological inference of conflict. We next show alignment length accounts for the high influence of two genes reported in empirical datasets. Finally, we also reiterate the impact misidentified orthology and short alignments have on likelihoods in large scale phylogenetics. We suggest that researchers should systematically investigate and describe the source of influential genes, as opposed to discarding them as outliers. Disentangling whether analytical or biological factors are the source of outliers will help uncover new patterns and processes that are shaping the Tree of Life.

## Introduction

It has become common practice to dissect the support individual sites and genes exhibit in favor of one topology over another (Lee and Hugall, 2003; Castoe et al. 2009; Smith et al. 2011). By identifying highly influential loci, researchers have shown that contentious relationships are sometimes the result of phylogenetic signal emanating from a small subset of the data (Castoe et al. 2009; Evans et al. 2010; Shen et al. 2017; Brown et al. 2017; Walker et al. 2018). These subsets may conflict with the majority of the data as a result of biological processes or analytical error. For example, a small number of sites overpowering the majority of signal in a mitochondrial dataset was reported by Castoe et al. (2009) as evidence for convergent evolution. However, in another mitochondrial dataset this was reported as inaccuracies in the modeling process (Evans et al. 2010). The identification of conflicting loci/sites with a large impact on species tree analyses or likelihood values has led to filtering practices. In analytical cases, where error in dataset assembly contributes to the conflict, filtering is justified. However, in cases where the strong signal of sites or loci (hereafter referred to as genes for simplicity) has a biological origin, filtering can lead to removal of informative (and sometimes the most reliable) data.

The difference among the likelihood of genes (the sum of the likelihood contributed by individual sites) when inferring one topology over another topology (influence) has been analyzed using Bayes factors, gene-wise likelihoods and/or partitioned coalescence support (Shen et al. 2017; Brown et al. 2017; Walker et al. 2018; Gatesy et al. 2019). The genes contributing the greatest influence, sometimes referred to as “outliers”, have been treated in a variety of ways throughout the literature. Outlier genes have appeared in phylogenomic analyses broadly across the Tree of Life. These outliers have been used as a basis for excluding genes from analyses (Nikolov et al. 2019), identifying errors in orthology (Brown and Thomson, 2017; Springer and Gatesy, 2018), analyzing conflicting signal among genes (Shen et al. 2017; Walker et al. 2018), testing for biological associations (Wang et al. 2019), and assessing the robustness of species tree hypotheses (Steenwyk et al. 2019). However, these analyses capture a mixture of biological and analytical signal, and this is why it is now important to dissect the source of the likelihood sum.

The topologies inferred using concatenation or coalescent analyses often disagree for historically contentious relationships. An emerging view is that these relationships will have the greatest amount of conflicting signal among genes and the removal of a small number of genes with the greatest likelihood sum often changes the relationships (Shen et al. 2017; Brown et al. 2017; Walker et al. 2018). Biologically speaking, these relationships may have been influenced by Incomplete Lineage Sorting (ILS), ancient hybridization, and other processes that make the histories of genes differ from the histories of the lineages that contain them. With high degrees of conflict, these relationships may, in fact, not be possible to resolve even using genomic datasets. However, by dissecting the properties of phylogenomic datasets, insight may be provided into why these relationships remain recalcitrant (Rokas and Carroll, 2006).

Non-biological forms of systematic and stochastic error also contribute to gene tree conflict, and these can mimic patterns generated by biological conflict (Richards et al., 2018). Current methods and models are often unable to extract the information from a single gene necessary to inform ancient divergences (Salichos and Rokas, 2013), with evolutionary phenomena such as saturation of sites removing information at deep time scales (Phillipe et al., 1994). Advances in modeling may eventually alleviate these issues, however, dataset curation also underlies some observed patterns (Springer and Gatesy, 2018). With small partitions of data having profound influence on inferred species relationships, which often form the basis of evolutionary studies, it is important to ensure all possible sources of error are accounted for. By disentangling the sources of signal, we can gain insight into not only what the inferred species relationships are, but also address the equally important question of *why* these relationships are inferred. Furthermore, detailed analyses provide insight into both the analytical and biological aspects of phylogenomic analyses.

Here we describe four specific reasons genes in a supermatrix may exhibit outlier behavior and the role this plays in inference, namely rate of molecular evolution, alignment length, misidentified orthology, and errors in modeling. Using both simulated and empirical examples selected across the Tree of Life, we show how these four factors can have a profound influence on phylogenomic analysis. We hope that these demonstrations will help researchers interrogate potential sources of error in their phylogenomic datasets and guide how analyses of gene influence in a matrix should be treated in future studies.

## Materials and Methods

### Simulation examples

All data and scripts have been deposited on Github (https://github.com/jfwalker/AnalyzingOutliers). All genes were simulated using seq-gen v1.3.4 (Rambaut and Grass, 1997) with the GTR model rate of molecular evolution, alignment length, misidentified orthology, and errors in modeling (transition rates of 1, 2, 3, 4, 5 and 6 and default state frequencies) and GAMMA rate variation with an alpha shape of 1.0 and divided into four categories. The length of the genes varied depending on the simulation used. The two-topology test was conducted by concatenating the simulated genes using the phyx v.99 program pxcat (Brown et al. 2017) and the site-specific log-likelihood (SSLL) values were calculated using RAxMLv.8.2.11 (Stamatakis, 2014) with “-f G” and the GTRGAMMA model of evolution. To obtain the gene-wise log-likelihoods (GWLL), we summed the log-likelihood of individual sites for each gene. To correct for length of the genes, the ΔGWLL (difference in GWLL between topologies) was divided by the length of the gene to obtain the average change in site-specific log-likelihoods (ΔSSLL).

#### Testing the effects of alignment length

To simulate the effects of alignment length, a birth-death tree was simulated using the phyx v.99 program pxbdsim (Brown et al. 2017) for five extant taxa. This was then scaled to a factor of one using the phyx v.99 program pxtscale (Brown et al. 2017). Simulations and inference of GWLL and SSLL were conducted for 1000 replicates. All genes were simulated on the same five-taxon topology (Supplementary Figure 1A), with four genes being 500bp in length and one gene 5000bp. The two-topology test was conducted using the topology upon which the simulations were performed and an alternative topology with one branch in conflict (Supplementary Figure 1B).

#### Measuring distribution of parameter estimates

Using the same topology as the one the alignment simulations were conducted on (Supplementary Figure 1A), we simulated two sets of 100 gene supermatrices. In the first set, 99 genes were 1000bp and one gene was 10bp in length. In the second set, all genes were 1000bp in length. A phylogenetic tree was inferred using maximum likelihood as implemented in IQtree v1.6.11 (Nguyen et al., 2015), with all genes partitioned “-q” and assigned individual GTR+G models of evolution. Parameter estimates were taken from the 10bp gene and the first 1000bp gene for the first and the second dataset respectively. The individual likelihoods for each partition were extracted using the “-wpl” option and divided by 10 or 1000 to obtain the average SSLL for the first and second dataset, respectively.

#### Simulation of misidentified orthology

To simulate misidentified orthology, we used a 12-taxon tree separated into two clades to simulate a duplication where each taxon retained a copy (Supplementary Figure 2A). The branch subtending the duplication was set at 0.1 subs/bp, 0.3 subs/bp, or 0.6 subs/bp and 1000 simulations of gene alignments were generated for each setting. In each simulation, all five genes simulated were all 1000bp in length. Orthology was “correctly inferred” by taking all the sequences from a 6-taxon duplicate (Supplementary Figure 2B), and incorrectly inferred by taking a mixture of sequences between the paralogous groups as illustrated in Supplementary Figure 2C.

#### Testing the effects of heterotachy

One thousand simulations were performed where four genes of 1000bp were simulated on a topology where the branch subtending the conflicting clade (Supplementary Figure 3A) was one fifth (0.1subs/bp) the length of the conflicting topology (Supplementary Figure 3B) upon which the fifth gene was simulated. The two topologies upon which the simulations were conducted were used for the two-topology test.

### Empirical Examples

The data for empirical examples from Walker et al. (2018), consisting of data from Chiari et al. (2012) and Walker et al. (2017), were downloaded from (https://github.com/jfwalker/MGWE). The ΔGWLL and average ΔSSLL values were calculated using the same script as the simulation examples. The empirical examples from Shen et al. (2017), coming from Wickett et al. (2014), were downloaded from (https://doi.org/10.6084/m9.figshare.3792189.v3). The gene-wise likelihoods were converted to average site-wise likelihoods using the partition file from a concatenation of the alignments made from pxcat, and the gene-wise likelihood inferred by Shen et al. (2017). The gene trees used were those from Shen et al. (2017), unless specified otherwise in the methods.

#### Testing for information content

Likelihood mapping was conducted using the quartet-puzzling approach (Strimmer and Von Haesler, 1997) as implemented in IQtree. This was done using the most influential per gene and per site alignments from Wickett et al. 2014 with regards to the monophyly of Bryophytes, and the third most influential gene of Chiari et al. 2012. This method allows an alignment to be assessed for its ability to resolve phylogenetic relationships by distinguishing how well quartets are capable of differentiating among conflicting alternative topologies.

The tree certainty (TC) scores (Salichos and Rokas, 2013) were obtained by inferring 100 regular bootstraps using IQtree with the GTR+G model of evolution. The TC score was obtained using the “-f i” option in RAxMLv8.2.12. The TC score is a sum of the IC (internode certainty) scores which ranges from −1.0 to 1.0 and can be calculated for all internal branches. Thus, a perfect TC score is *n-3* for an unrooted tree, where *n* is the number of taxa. In short, the IC score, which forms the basis of the TC score, is calculated by testing how many clades within a set of BS trees conflict with the ML tree and how those conflicts relate to one another. Assuming all BS trees are identical to the ML tree, the IC value is 1.0, and assuming all BS trees support the same alternative topology the IC score is −1.0. Thus, if there is no information for resolving the relationships and the resolution for a given relationship amongst the BS trees is completely random the value is 0.

#### Orthology analysis of the vertebrate dataset

To examine if another gene may have potential issues in orthology detection from the vertebrate dataset, we downloaded the nucleotide sequences for the chicken genome (GRCg6a) from ENSEMBL (download date: 8/21/2019). A BLASTN database was generated from the chicken genome and the third most influential gene from Chiari et al. 2012 (ENSGALG00000008314) was used as a query against the genome database, with an e-value setting of 1e-3. All matches were combined and aligned using PRANK (Löytynoja and Goldman, 2008) with the -codon option turned on. A phylogenetic tree was inferred using maximum likelihood as implemented in IQtree v.1.6.11 (Nguyen et al., 2015), with the GTR+G model of evolution with 1000 ultrafast BS (Hoang et al. 2017).

#### Comparison of ML topology conflict vs. ΔGWLL conflict in the carnivorous plant order Caryophyllales

The gene trees from Walker et al. (2017) whose edge matched either the inferred Astral (Mirarab et al. 2014) topology or the inferred ML concatenation topology from the original study were identified using PhypartsPy (https://github.com/jfwalker/CompMethodsCode). The program identifies edges on rooted trees between two topologies and reports the concordant/conflicting edges, the support values for each edge, and the length (subs/bp) of the edge. The Astral topology contained an edge consisting of *Ancistrocladus robertsonorium* (MJM2940), *Drosophyllum lusitanicum* (DrolusSFB), *Nepenthes alata* (NepSFB), and *Nepenthes ampullaria* (Neam), which is in conflict to the supermatrix inferred edge of *N. alata, N. ampullaria, Aldrovanda vesiculosa* (MJM1652), *Dionaea muscipula* (Dino), and *Drosera binata* (DrobinSFB). The ΔGWLL was compared to the total number of substitutions inferred along a given branch, calculated by multiplying the edge length (subs/bp) by the alignment length (bp), thereby generating a value for the estimated number of substitutions that occurred along the branch.

The likelihood of the gene (named cluster3522 in the original study from Walker et al. (2017)) was chosen to be evaluated because the topology of the gene tree was identical to the Astral tree, however, the ΔGWLL indicated support for the supermatrix tree from the original study. To ensure a fair comparison among likelihoods, branch lengths were re-estimated on both the Astral and the Supermatrix topology using IQ-TREE with the “-q” option and the GTR+G model of evolution was applied to all partitions. The gene tree was re-inferred with IQ-TREE and the GTR+G model of evolution to ensure the topology did not change as a result of the likelihood inference method or program. The likelihood was then evaluated for all three sets of branches using IQ-TREE with the “-show-lh” option and branch length parameters fixed.

## Results and Discussion

The use of large datasets in statistical phylogenetics has shown that specific relationships in the inferred maximum likelihood topology (ML) may be influenced by less than one percent of sites or genes (Lee and Hugall, 2003; Castoe et al. 2009; Evans et al. 2010; Brown and Thomson, 2017; Shen et al. 2017; Walker et al. 2018). Although it should be expected that not all sites and genes (sum of site likelihoods for a partition) contribute the same value to the final likelihood score, understanding why some genes appear to exhibit outlier behavior, with differences in likelihoods for alternative topologies hundreds of times greater than others, is important for understanding how the Tree of Life is shaped. In the following sections we discuss four common biological and analytical (i.e., non-biological) reasons genes exhibit outlier behavior. Specifically, we use simulations and empirical data selected across the Tree of Life to demonstrate how genes can exert drastically different influence, support for one topology over another, hereafter called the ΔGWLL. We discuss how the ΔGWLL has been used in the literature to analyze conflict, filter gene, and identify errors in orthology or in modeling. Furthermore, we extend the discussion to explain some ways this can be used to disentangle biological and analytical sources of influence. We hope this continues to build on the discussions of how to proceed using large scale phylogenetic datasets.

### Inferred molecular rate can mislead conflict analyses

The distribution of conflict allows researchers to begin modeling evolutionary events, however, current models and methods assume that the conflict is biological. In this section we explore the effect molecular rate has on the influence of a given gene. We also show how molecular rate is capable of generating gene trees whose inferred maximum likelihood topology conflicts with the topology the gene supports in a ΔGWLL analysis.

#### Simulating on an induced molecular rate shift generates disproportionate influence

From an analytical standpoint, the molecular rate of evolution is a proxy for the amount of information a gene contributes to a supermatrix analysis. If a gene has a disproportionate rate of evolution, then the gene will likely have a greater influence on the inferred topology. Examining tree length (total inferred subs/bp) can help identify this, but if the question is regarding a specific relationship then the length of the edge subtending the relationship of interest is an important factor to consider. Through simulation it is possible to isolate one edge to check the influence a rate shift has on inference. If most genes support one relationship, while a small number of genes support a conflicting topology with a major rate shift (Supplemental Figure 3), then this can have disproportionate consequences on a test between two relationships (Figure 1A).

**Figure 1.**
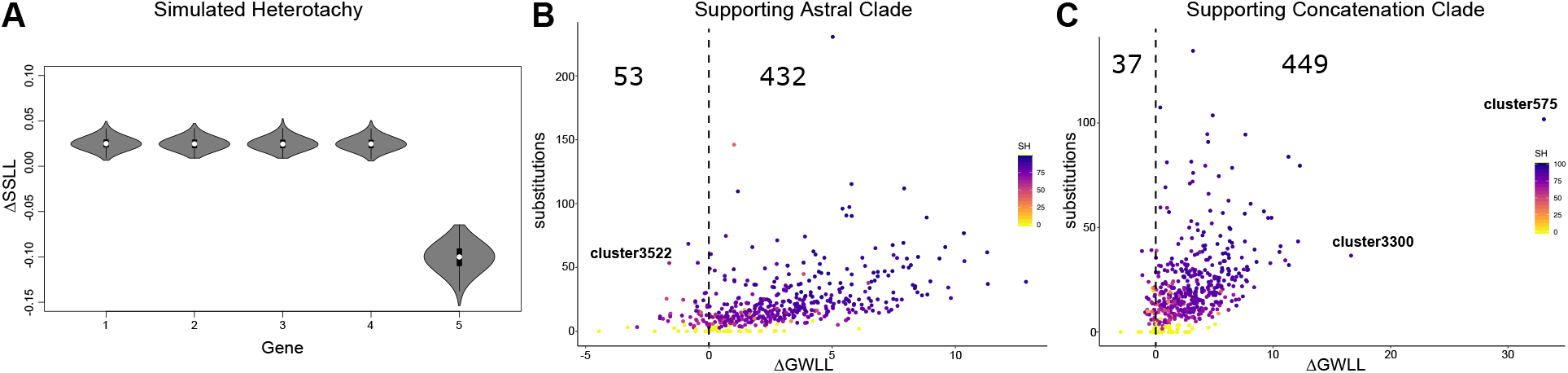
Molecular rate of evolution can result in altered gene influence and can change which topology is supported. A) Violin plots of 1000 simulations where the first four genes are simulated on the same topology and the fifth gene is simulated on a topology where a rate shift has taken place on the edge subtending the conflicting relationship. B&C) Plots of the ΔGWLL values between the topology of Walker et al. (2017) inferred by Astral vs. that inferred by concatenation. The y-axis depicts the substitution inferred to have occurred at the node subtending the contention relationship for each gene tree. Each point is colored based on the SH support from the original study. B) The 485 genes from the carnivory dataset whose ML tree contains the same clade as Astral. 432 genes whose ΔGWLL is greater than 0 support the Astral clade from a GWLL analysis, and 53 genes with ΔGWLL less than 0 support the concatenation topology from a GWLL analysis. The gene ‘cluster3522’, further explored in supplementary 4 is labeled. C) The 486 genes whose ML tree topology contains the same contentious clade as the concatenation topology. 449 genes with a positive value support their ML topology in a ΔGWLL framework and 37 genes with a negative value support the Astral topology over their ML topology in a GWLL framework.

#### The maximum likelihood gene tree and the ΔGWLL support different species trees in ten percent of tested cases

To analyze the influence of molecular rate on empirical data we identified all genes from the study by Walker et al. (2017) whose maximum likelihood gene tree topology matches the Astral topology in the study (485 genes). We examined both the relationship of the ΔGWLL to the molecular rate and how many of these genes support the Astral topology based on the ΔGWLL two-topology test (Figure 1B). We found that of the 485 genes whose maximum likelihood (ML) gene tree supports the Astral topology, 53 of them support the concatenation topology when analyzing conflict on the basis of ΔGWLL values. Although it is counterintuitive that an ML gene tree and the ΔGWLL would support different topological resolutions, this arises by the model evaluation process. The branch length parameters of both the concatenation and the Astral topology are inferred using all genes in the dataset, thus they are likely suboptimal for the gene tree. As a result, the sub-optimal model inferred by Astral is a better fit than the sub-optimal model inferred by the concatenation. This is similar to the 266 genes within the Walker et al. (2017) dataset, whose ML topology did not contain either the Astral or the concatenation clade. In the ΔGWLL measure of conflict those genes were all forced to support one suboptimal topology over another.

To examine this in more detail, we investigated cluster3522 which has an inferred gene tree topology concordant with the supermatrix, however, within the ΔGWLL framework cluster3522 supports the Astral topology (Figure 1C). To further examine this phenomenon, we inferred the log-likelihood of three models, the supermatrix ML topology model (concatenation), the Astral topology model, and the gene tree ML topology model. The log-likelihood of cluster3522 evaluated on the ML gene tree for the gene was −36,091, whereas the log-likelihood evaluated on the concatenation topology was −36,265, and the log-likelihood on the Astral topology was – 36,267 (Supplementary Figure 4). This means that when evaluated under the sub-optimal parameter estimates that come from the concatenation framework, cluster3522 has a better fit to the model where the Astral inferred contentious relationship is. This highlights that conflict evaluated using ΔGWLL, can be an evaluation of conflict under suboptimal parameter estimates and is thus prone to supporting a topology different from its own ML.

Evaluating conflict on the basis of ΔGWLL provides important information into how the genes interact to infer the species tree and should thus be explored. Looking at the numbers of genes that support either species relationship is informative as to why the relationship was inferred, but is not the same as evaluating conflict among ML trees. These two measures provide complementary but different information. This is important to consider before inferring biological processes from conflict analysis done using ΔGWLL.

Rate heterogeneity helps contribute to this, because even if the gene topology matches the most likely topology, different rate parameters (branch lengths) can cause a better fit to an alternative topology. This rate difference also likely explains the superior AIC score that appears when all branches are estimated separately (Walker et al. 2018) as opposed to being edge-equal (i.e., equal across partitions), because in the example factoring in the heterotachy of cluster3522 the likelihood within the supermatrix would be the same as that of the ML tree. Accounting for heterogeneity in rates among genes through the edge-proportional branch length model in a supermatrix analysis has been shown to recover the same result as summary methods (Duchene et al. 2018). This could be explained by some coalescent summary methods being based on topology and therefore not influenced by disparate likelihoods or molecular branch lengths. As the field continues to develop, it will be important to validate methods capable of accounting for inter gene rate and topological variation.

#### Biological implications and influence on understanding the Tree of Life

Molecular rate heterogeneity is common among genes and has long been used as a test for a gene to be under selection. Supermatrix methods are especially prone to being affected by this, and genes that coalesce deep in the tree should theoretically have longer branches. Supplementary Figure 3 could be seen as an extreme example of deep coalescence, where four genes show one relationship and the fifth has the most influence as it has experienced the greatest amount of change (Figure 1A). Understanding the source of rate variation is thus key to understanding the Tree of Life and potentially key to understanding the reliability of a gene to speak to any given relationship.

Although it may be difficult to know the underlying cause, it is imperative to analyze genes exhibiting such behavior in detail and work to understand why the ΔGWLL’s originate. As the number of phylogenomic studies examining likelihood signal increases, it will be possible to test for the emergence of patterns on larger scales. Such analyses may reveal whether the influence of a gene is the result of shared common properties (e.g., more likely to arise in large multigenic families) or functions (e.g., more likely to code for enzymes from certain metabolic pathways) or not. This should improve our ability to predict and account for disparate influence and, by examining the role such genes fulfill in organisms’ development and function, it may connect the level of influence to biology.

### Alignment length may underlie highly influential genes

#### Analytical sources of influence

One explanation for disparity in the likelihoods and outlier behavior is that genes often have different alignment lengths. Assuming all sites in a statistically consistent data matrix have the same likelihood, the likelihood value calculated for a gene will be correlated with alignment length, and therefore the most influential gene would be the longest gene. A 5000 bp gene region will be more influential than a 500 bp gene region (Figure 2A). Correcting for alignment length (Figure 2B) provides a measure of the average contribution of each site to signal.

**Figure 2.**
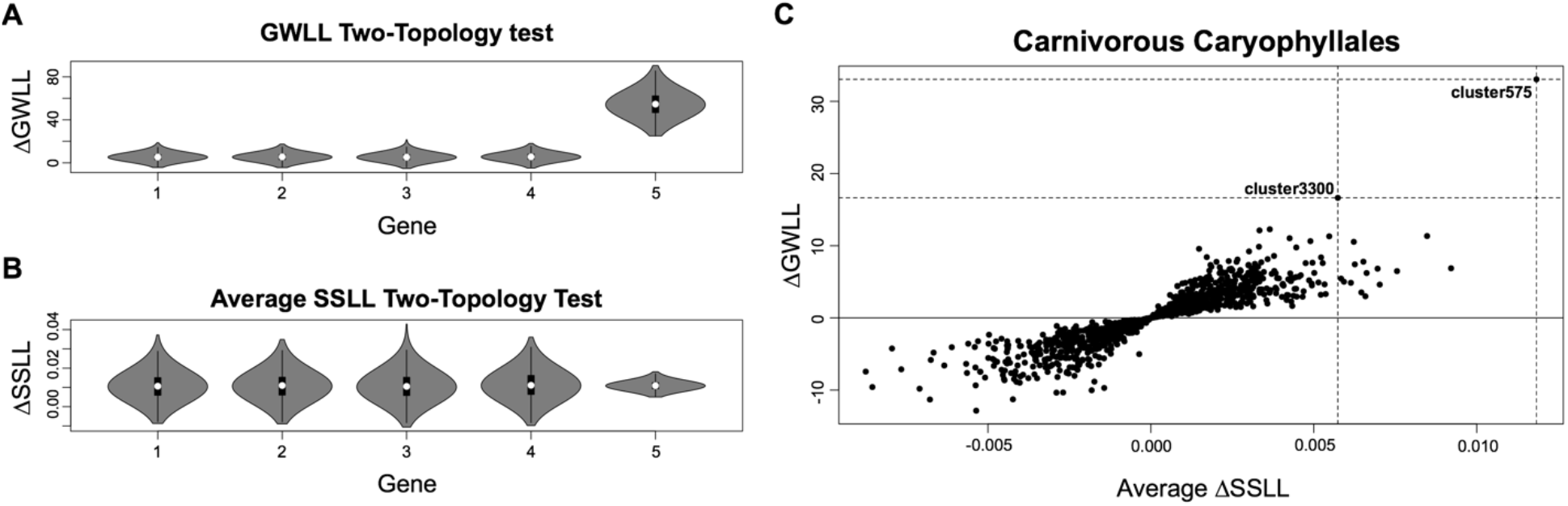
Adjusting for alignment length helps uncover a source of outlier behavior. A&B are violin plots for 1000 simulations of alignment lengths influence on ΔGWLL and ΔSSLL. A) Shows the change in ΔGWLL with gene 5 (5000bp) exhibiting significantly higher loglikelihood density than the other four (1000bp). B) Shows the ΔGWLL divided by the length of the alignment to obtain the average ΔSSLL. C) Shows an empirical example from Walker et al. 2017. The two genes identified as “outlier genes” in the original study are labeled. The horizontal lines are used to mark the point where the topology the gene supports switches 0 (solid line), and the ΔGWLL of the outlier genes (horizontal dashed lines). Vertical dashed lines are placed on the x-axis to mark the average ΔSSLL of the outlier genes.

#### Outlier behavior may be explained by alignment length in an empirical dataset

We revisited the two most influential genes in the Walker et al. (2017) carnivorous Caryophyllales dataset. These genes were termed “outlier genes” as their removal altered the species tree and they had already been examined to find no identifiable modeling issues or errors in orthology detection (Walker et al. 2018). When adjusting for alignment length, one of the outlier genes (cluster3300), previously the second most influential gene from the ΔGWLL analysis, became the 16^th^ most influential gene among those supporting the supermatrix topology (Figure 2C). This gene is 2,900 base pairs long, while the average gene length of the dataset is 1,711. Thus, the disproportionate influence of cluster3300 in the supermatrix is, at least in part, a result of disproportionate length. As with any empirical example, the disparate signal is unlikely to be isolated to a single factor, however, the longer alignments have a greater potential to gain influence from either having a cumulative effect of summing the differences in likelihood or containing subsets of highly influential sites. The most influential gene from Walker et al. (2017) (cluster575), was also the most influential gene per site (Figure 2C). Thus, the disproportionate influence of cluster575, which was also shown to not be attributed to model violation or orthology error, cannot solely be attributed to alignment length which was 2,793bp.

#### Biological processes and influence on understanding the Tree of Life

There is natural variation among the length of genes. Therefore, in the age of phylogenomics when sequences are no longer limited to the length of a PCR reaction, the Tree of Life is likely becoming skewed towards the evolutionary relationships inferred by longer genes. Nevertheless, the greater influence of longer alignments is not intrinsically *bad*, it is the result of the sites being assumed to be completely independent and identically distributed, so likelihood is the cumulative score across sites. The greater length indicates more data that may correspond to more useful signal to estimate the parameters in statistical phylogenetics.

### Misidentified orthology may positively mislead studies

#### Analytical origin

Misidentified orthology has previously been identified as the source of disparate signal from the most influential genes (Brown and Thomson, 2017). The importance of accurate orthology inference is imperative and thoroughly discussed in the literature (Eisen, 1998; Gabaldón, 2008; Salichos and Rokas, 2011; Springer and Gatesy, 2018). Orthology inference is essential for accurate phylogenetic reconstruction: as orthologs capture the speciation event, misidentified orthology thereby captures a duplication event. The penalty of misidentified orthology in a supermatrix can result in a greater number of inferred substitutions between two taxa. If a gene alignment is, in fact, not comprised of orthologous sequences, the phylogenetic relationships reconstructed will be misleading and there will likely be a long edge subtending the taxa derived from different orthologs and thereby generate greater signal (Supplementary Figure 2; Figure 3A).

**Figure 3.**
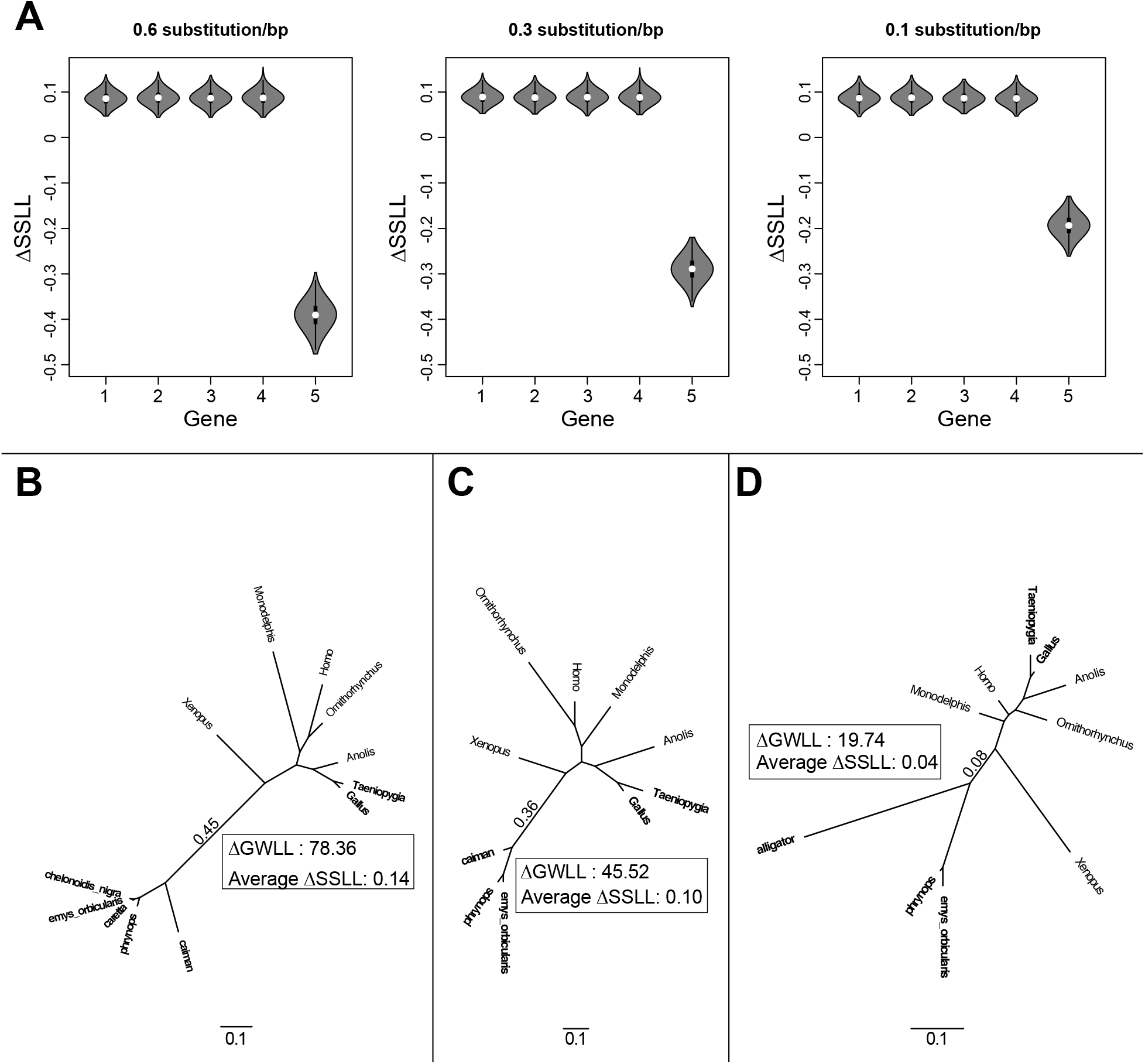
Errors in orthology detection result in disparate signal whose magnitude increases with the branch subtending the misidentified ortholog. A) Violin plots depicting the 1000 replicates of simulated data where the first four genes had correctly identified orthology and the next gene was incorrectly identified (Supplementary Figure 3). Decreasing the length of the branch subtending the orthology in conflict led to a decrease in magnitude of the signal. B-D) Incorrectly identified orthologs from Chiari et al. (2012) presented in decreasing order of their influence, where the branch subtending the conflict has the length in subs/bp labeled, with the GWLL and SSLL values from the two-topology analysis labeled. Bolded taxa are those regarding the contentious relationship B) is gene 8916, C) is gene 11434 and D) gene 8314.

This type of error arises due to the increased number of substitutions that separate the clade in question from the other possible relationships. This increased number of substitutions can also be the underlying cause of the misidentified orthology (Smith and Pease, 2016), as similarity searches, such as blast, will minimize difference in substitutions, but will not account for covariance of the substitutions. The likelihood calculated may reflect the evolutionary history of the sequences in question, however, in the case of misidentified orthology, this history will not be reflective of the species history.

#### Empirical Examples

When accounting for alignment length while analyzing the Chiari et al. (2012) dataset the two genes (8916 and 11434) previously identified by Brown and Thomson, (2017) as the most influential remained so. We then examined the third most influential gene (8314), as this also remained highly influential when accounting for alignment length. The alignment is 441bp, likelihood mapping showed only 12.4% of the quartets were uninformative (Supplementary Figure 4C), and the TC score was 1.59 out of a possible 12. Taken together, these results indicate that this gene does not exhibit the same issues as the Bryophyte gene (7159_C12) and should have sufficient information to inform the model of molecular evolution. To first look for possible errors in orthology, we examined the evolutionary relationship of all sequences in the orthology group “ortholog” to all BLAST inferred homologs from the chicken genome. The inferred phylogeny had chicken genome homologs appearing in three separate places (Supplementary Figure 5). This indicates that those duplications are not all specific to chicken and that 8314 is likely composed of at least three separate orthologs.

We then examined the three orthology-based outlying genes to identify the properties that may drive this signal (Figure 3 B-D). This provides an empirical example where inferred number of subs/bp correlate with the level of influence through both the ΔGWLL and average ΔSSLL. When placed in a supermatrix framework the total number of substitutions would then be accounted for and the influence of the subs/bp would become a function of that and the alignment length at the edge. Although, in this empirical example it should be noted that complex other influences (e.g., taxon sampling) were not accounted for.

#### Biological implications and influence on understanding the Tree of Life

Although patterns of conflict found in predicted orthologs such as those seen in the Chiari et al. (2012) dataset may appear to have arisen by analytical error, similar patterns may arise through biological processes as well. The process of gene duplication and loss, something especially prevalent in plants, has the potential to give rise to the same signal, as true orthologs become lost, and only the paralogs may be recovered from the genome. This pattern may also emerge as the result of horizontal gene transfer. This can mimic misidentified orthology and has the potential to be an under-appreciated source of conflict (Dunning et al. 2019). Through data interrogation, the biological processes that generate this form of signal are likely to emerge and help gain a clearer understanding of the Tree of Life.

### Model misspecification and lack of information increase variance-

#### Analytical source of influence

Information and alignment length should be tightly correlated as both invariable and variable sites contribute to the signal. Lacking information or not having a proper model available can all contribute to poor model fit. Although quantifying the exact amount of information in an alignment is a difficult task (Strimmer and Von Haesler, 1997; Goldman, 1998; Townsend, 2007), the amount of information plays a crucial role in the ability to infer evolutionary relationships. Similar to most statistical problems it is rarely advisable to estimate more parameters than you have data (Burnham and Anderson, 2002), however, genes are finite in length and, it may not be possible to accumulate more data for the alignment. Current methods are being developed to address this issue (Benoit et al. 2020), however, until these become a mainstream part of analytical pipelines, the ML tree inferred from fewer data points than parameters may be positively misleading.

In statistical phylogenetics the tree itself is part of the model, whose topology acts as a constraint during the tree search, and therefore the inferred topology is dependent upon the parameters which compose the tree. A gene that is run under the GTR+G model of evolution would have five parameters from the transition rate matrix, three from the estimated base frequencies, and one contributed by estimating the alpha parameters of the gamma distribution. A further 2*n*-3 (where *n* is the number of taxa) parameters are estimated in an unrooted phylogeny. As demonstrated through simulations on 10 bp vs. 1000 bp data, in a supermatrix where all other sequences are 1000 bp, the variance among parameter estimates is still far greater using 10bp (Figure 4), and this results in greater variance of the estimated log-likelihood scores for that segment. In a single gene phylogenetic analysis this variance should be captured by the bootstrap, but as part of the larger matrix the bootstrap is no longer able to capture this variance (Seo, 2008). Although this represents a very trivial simulation, and the random generation of substitutions through the simulator may explain the results, a lack of data is a common aspect of large-scale phylogenies.

**Figure 4.**
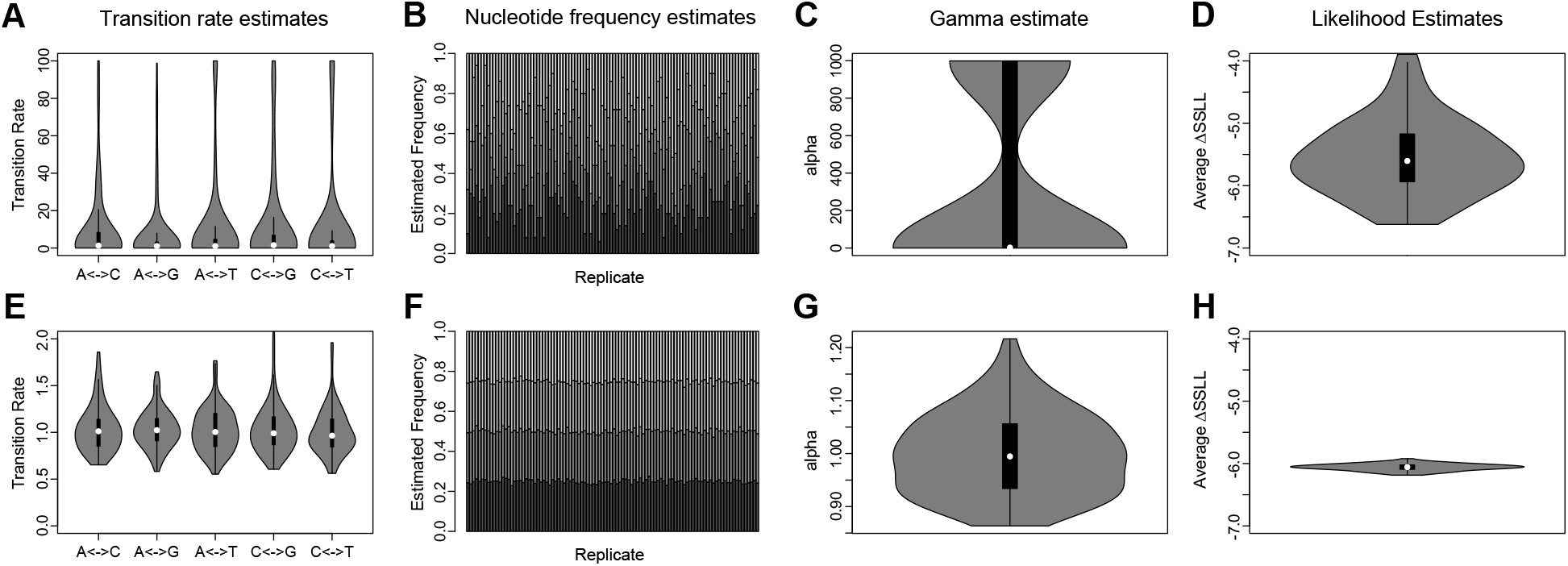
Shorter sequences generate a greater variance in parameter estimates and average ΔSSLL. A-D) Parameter and log-likelihood estimates for a 10bp sequence in a 100 gene supermatrix, where the other 99 genes were 1000bp in length. E-H) Parameter and log-likelihood estimates for a 1000bp sequence in a supermatrix where all sequences were 1000bp in length. A&E) Violin plots depicting the distribution of estimated transition rates. All simulations were conducted using JC and thus the true rate for each transition is 1.0. B&F) Estimated nucleotide frequencies, each shade depicts a difference nucleotide and the true frequencies are 0.25 based on the 100 simulations. C&G) Violin plots depicting the distribution of the shape parameter alpha values estimated for the gamma distribution. D&H) Violin plots depicting the distribution of average ΔSSLL.

#### Empirical Examples

We analyzed the average ΔGWLL in comparison to average ΔSSLL for the most influential genes from Chiari et al. (2012) and Wickett et al. (2014), identified by Brown and Thomson, (2017) and Shen et al. (2017) respectively. The bryophyte comparison provided insight into how modeling and information may influence signal (Figure 3). Although, this may not be representative of the behavior of the gene in the supermatrix, analyses of the single gene provide insight into how the gene may contribute to the overall inferred tree.

The topology of the most influential gene by ΔGWLL (6349_C12) and 26^th^ most influential gene supporting monophyletic bryophytes by average ΔSSLL recapitulates many well-supported relationships in the land plant phylogeny (Figure 5A). This gene gains influence in part due to its alignment length, as it appears to lose notable influence when alignment is accounted for. However, the most influential gene by average ΔSSLL (7159_C12) does not recapitulate many well-supported relationships (Figure 5B). The gene itself contribute only 12bp, compared to 1049 bp of 6349_C12, in a dataset where the average gene length is 455 bp. Although the gene is situated in a supermatrix and thus testing the information it is capable of contributing in that framework is difficult, we tested how much information the gene itself contained through an analysis of the gene tree topology with TC scores and the alignment through likelihood mapping (Supplementary Figure 6).

**Figure 5.**
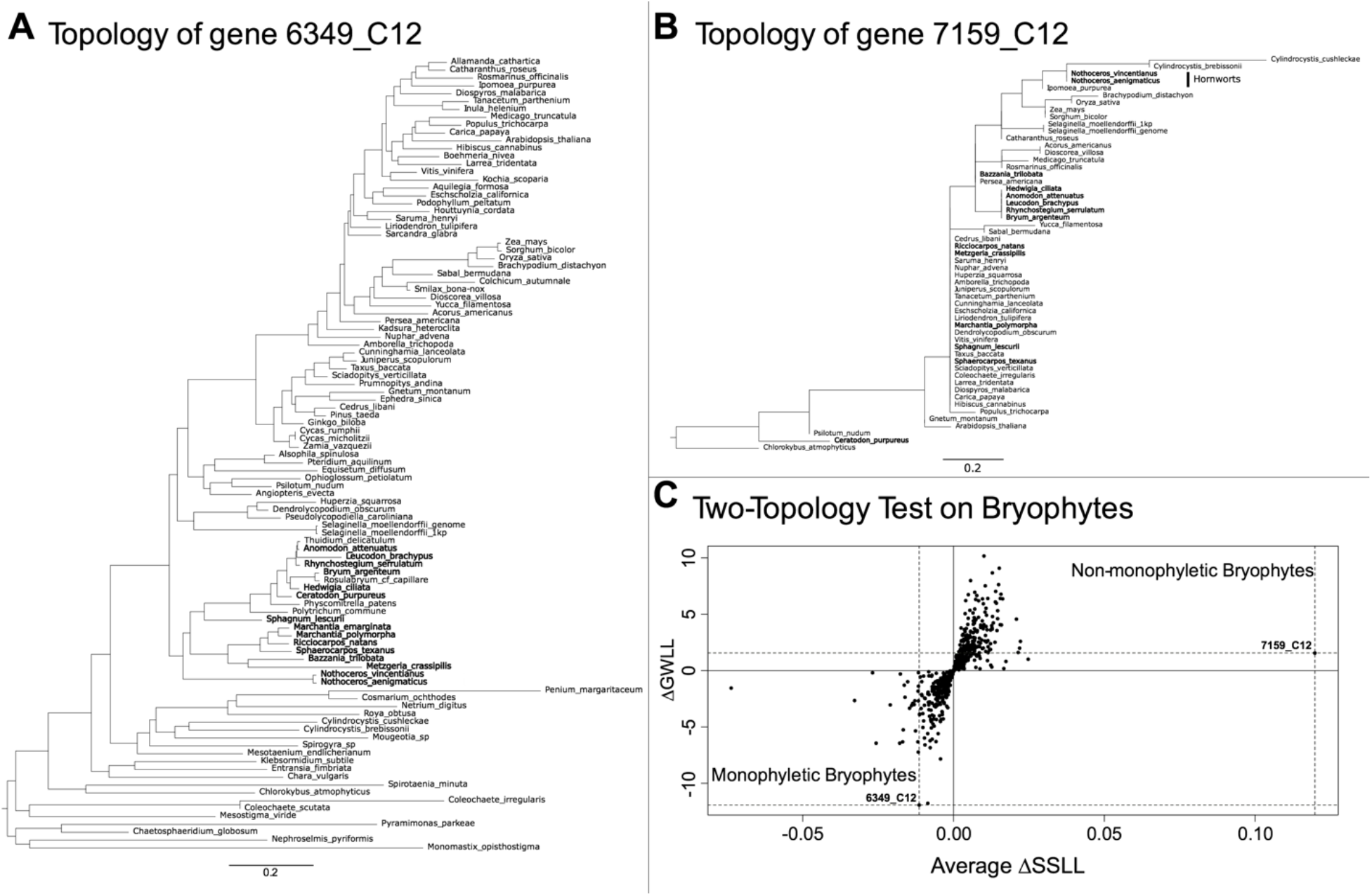
Examining the topologies of the most influential gene (highest ΔGWLL) and the most influential per site gene (highest Average ΔSSLL) from Wickett et al. 2014 gives insight into the source of their high influence. A&B) The highlighted names are species in which both gene 7169_C12 and 6349_C12 were found. A) The topology of 6349_C12, the gene with the greatest ΔGWLL. B) The topology of 7169_C12, the gene with the greatest average ΔSSLL. C) The average ΔSSLL vs. the ΔGWLL, solid lines mark the change in support from monophyletic bryophytes to non-monophyletic bryophytes. Dashed horizontal and vertical lines mark the genes with the highest ΔGWLL and highest average ΔSSLL.

We found that the TC score for the most influential gene by average ΔSSLL was −21 with a maximum possible score of 52. This indicates that within the bootstrap replicates there is a bias towards relationships that are not found in the ML gene tree. To analyze the information in the alignment we performed likelihood mapping and found that 70.6% of the quartets that compose the gene tree were uninformative (Supplementary Figure 6). Both these metrics would indicate that 7159_C12 lacks the necessary information to resolve the gene tree. The same two evaluations on gene 6349_C12 found that 27.5% quartets were uninformative (Sup Fig 4B) and the TC score was 33.34 out of a possible score of 98.

#### Biological Reasons and influence on understanding the Tree of Life

Molecular evolution is subject to far greater effects than what can be captured through models of evolution. It has been demonstrated that increasing complexity of the model, can in fact alleviate significant conflict (Beaulieu et al. 2018; Evangelista et al. 2019). Ensuring proper fit through model testing is undoubtedly valuable even in the era of large-scale phylogenetic datasets (Brown and Thomson, 2018), as it may help unmask one reason for these influential genes. Compositional heterogeneity across the Tree of Life should be considered (Foster, 2004), however, the vast majority of phylogenetic inference is conducted with only one model of molecular evolution used to estimate the entire topology.

## Conclusion

Although statistically inferred phylogenies depict evolutionary relationships, they are models, and like all evolutionary models, they cannot realistically be expected to capture all of the underlying biological complexity. This is not to say that the species relationships inferred are inaccurate, or that inferring them is not a worthwhile pursuit. Rather, as we reduce the uncertainty of what we are modeling we begin uncovering new and exciting evolutionary patterns. Methods such as Bayes Factors or ΔGWLL should continue to be investigated and expanded upon, because the interactions between the methods we use, and the data analyzed are complex and nearly impossible to account for all scenarios.

In this paper we have explored several common sources of disproportionate influence, however, with greater data dissection researchers continue to discover more (e.g., varying signal across chromosomes (Crowl et al. 2019; Li et al. 2019), or the influence of taxon sampling (Walker et al. 2017; Shen et al. 2018) and individual sites (Shen et al. 2017)). One approach to handling these is to avoid incorporating likelihood score as is done in some coalescent-based summary methods. However, differences in likelihood show valuable information about datasets and can help identify biological and analytical properties of the data.

Now that genomic data is readily available for many species, it is essential that we understand the intricacies of the data. Genome-scale data does not represent one tree, it represents a set of trees. Methods for accommodating this heterogeneity have undoubtedly brought us closer to the Tree of Life. Disparity in influence marks an area for exploration as this represents one of many complex properties of this data that remains to be uncovered, understood, and incorporated into phylogenetic and functional analyses.

## Acknowledgments and Funding Information

The authors would like to thank N Walker-Hale, S Walker, D Larson, JW Brown and J Chang for helpful comments on earlier drafts of this manuscript. This work was supported by the Gatsby Charitable Foundation (PTAG/022) and the Isaac Newton Trust/Wellcome Trust ISSF.

## Notes

https://github.com/jfwalker/AnalyzingOutliers

### Competing Interest Statement

The authors have declared no competing interest.

