## Supplementary Figures 1-6 for "Disentangling biological and analytical factors that give rise to outlier genes in phylogenomic matrices"

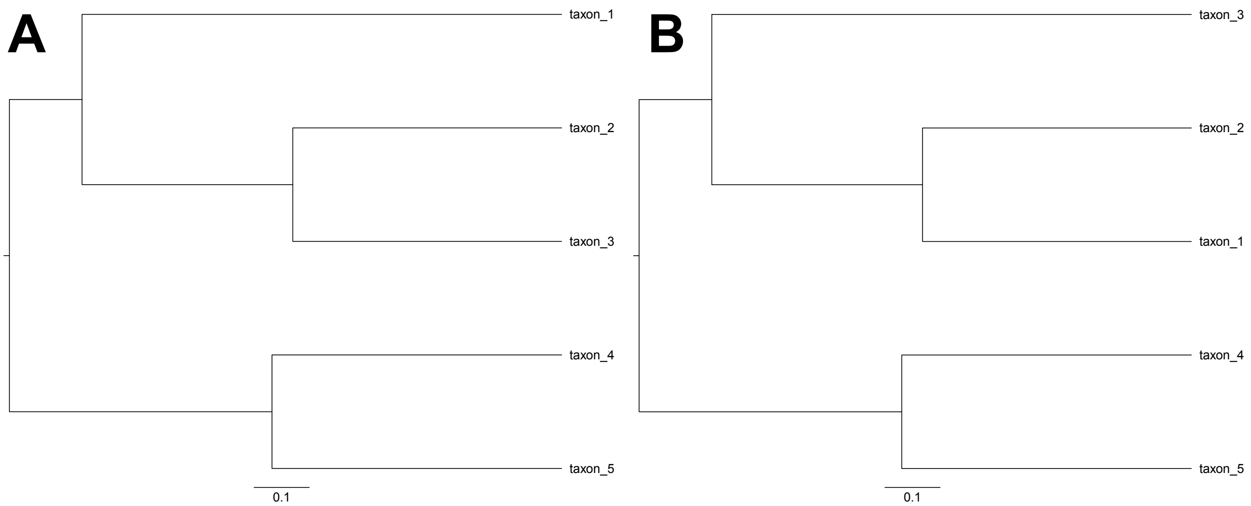


**Supplementary Figure 1. Topologies used for alignment length simulation.** A) Topology upon which all sequences were simulated. B) Alternative topology where the positions of taxon_1 and taxon_3 are switched. The SSLL values were calculated upon both topology A and B.

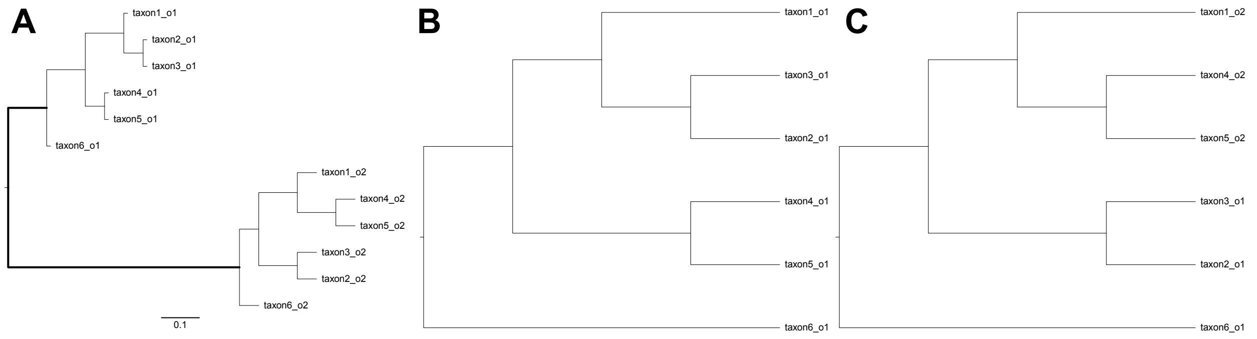


**Supplementary Figure 2. Topologies used for simulation of misidentified orthology and for the two-topology comparison.** A) The topology upon which simulations were conducted. The length of the highlighted branch was either 0.1, 0.3, or 0.6 subs/bp depending on the simulation. Sequences that end in "_o1" were used for the correctly identified orthology topology and the basis of topology 1 shown in B. Bolded sequences were used for the incorrectly identified orthology topology shown in C and as the fifth gene in the two-topology analysis. B&C) The two topologies used in the two-topology test.

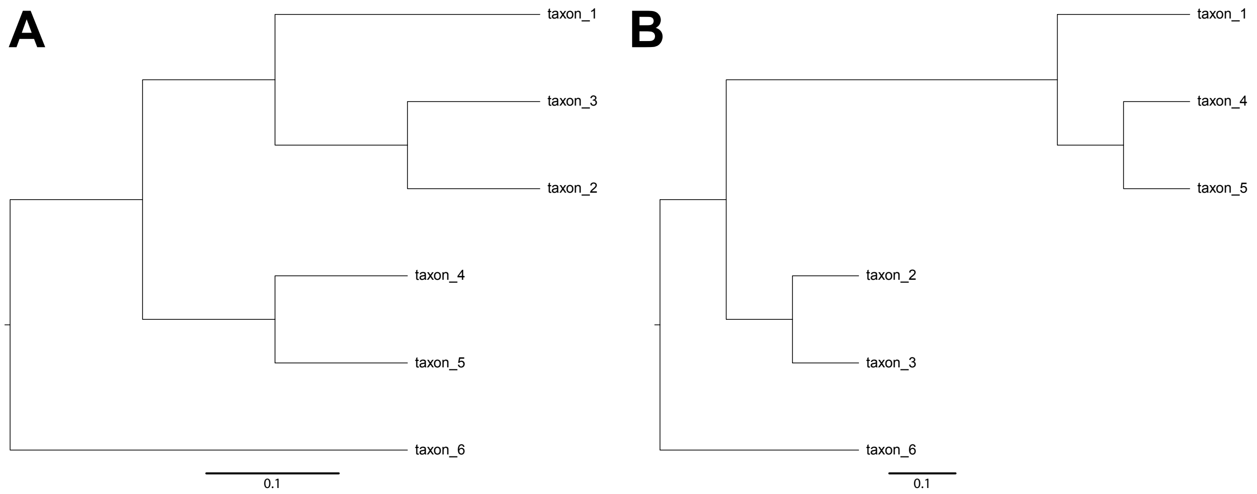


**Supplementary Figure 3. Topologies used in the heterotachy simulation.** A) Topology upon which the first four genes were simulated. B) Topology upon which the fifth gene was simulated where the branch subtending the conflicting clade has a 5x rate shift. The two-topology comparison was conducted using A compared to B, where branch lengths were re-estimated using the concatenated five gene supermatrix.

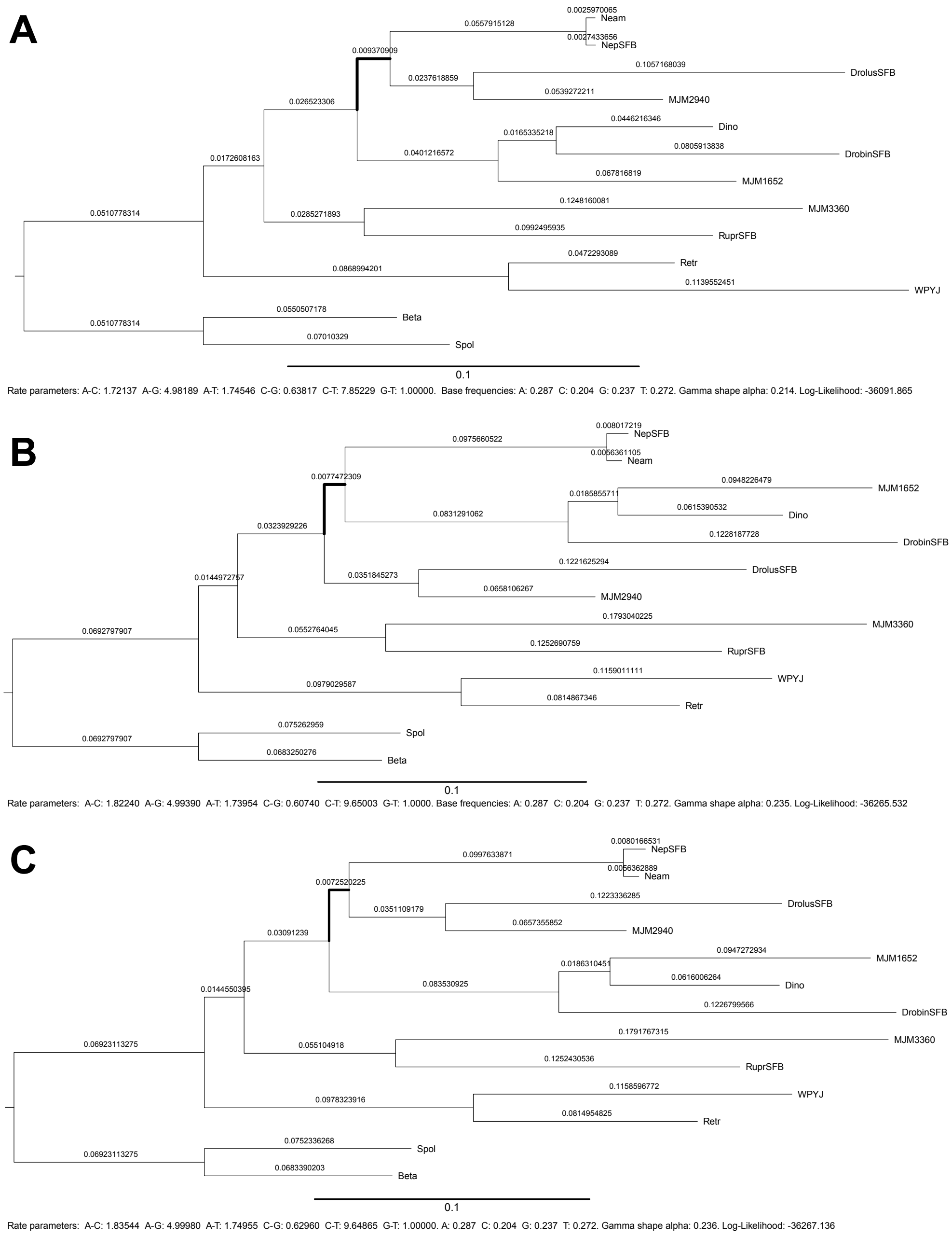


**Supplemental Figure 4. Likelihood analysis on sub-optimal parameters changes the inferred relationship of cluster3522.** The edge defining the contentious clade is highlighted on each topology and topologies are sorted based on likelihood. All parameters of the tree are depicted with the inferred log-likelihoods, trees are rooted, therefore the edge defining the root has been divided into two edges of equivalent length. A) The topology with all parameters maximized using only the sequence data from cluster3522. B) The topology inferred by concatenation, with parameters estimates of linked branches derived from all genes in the supermatrix and log-likelihood calculated across these parameters. C) The topology inferred by Astral, with parameters estimates of linked branches derived from all genes in the supermatrix and log-likelihood calculated across these parameters. The optimized likelihood and values for GTR and Gamma were obtained by evaluating the tree without optimizing branch lengths.

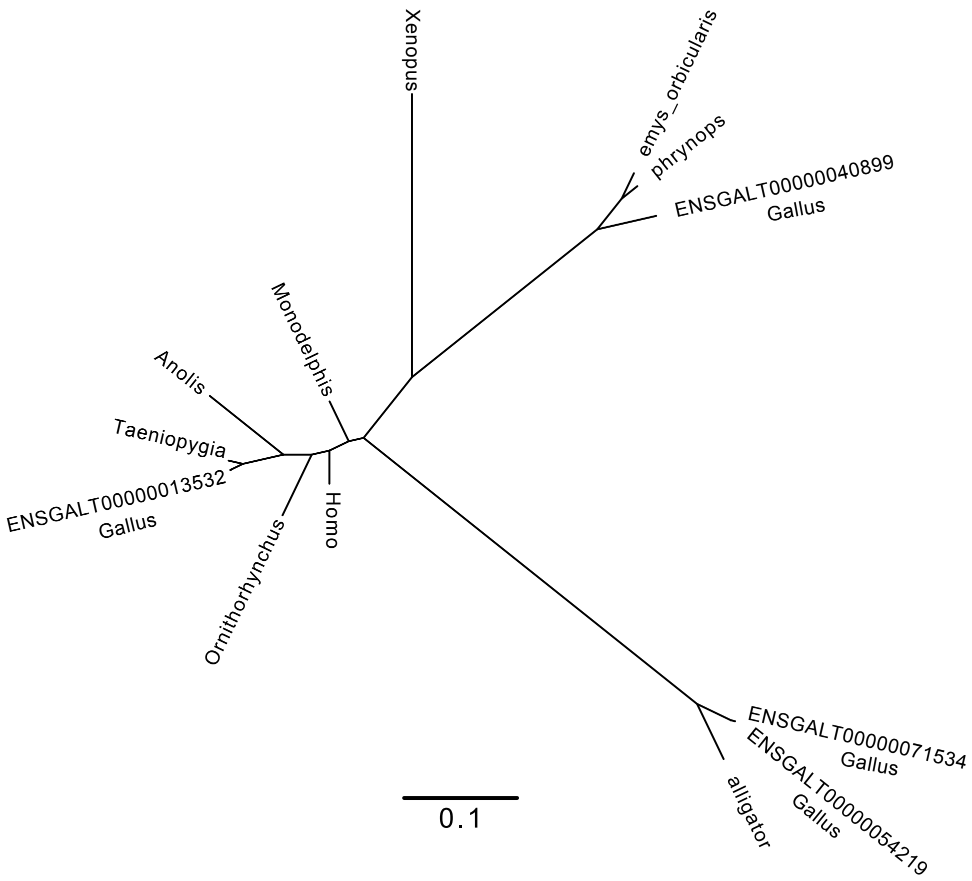


**Supplemental Figure 5. Inferred ortholog corresponding to 8314 from Chiari et al. (2012) clusters with three paralogs.** Inferred ML tree for gene 8314 from Chiari et al. (2012), with sequences for the *Gallus gallus* genome.

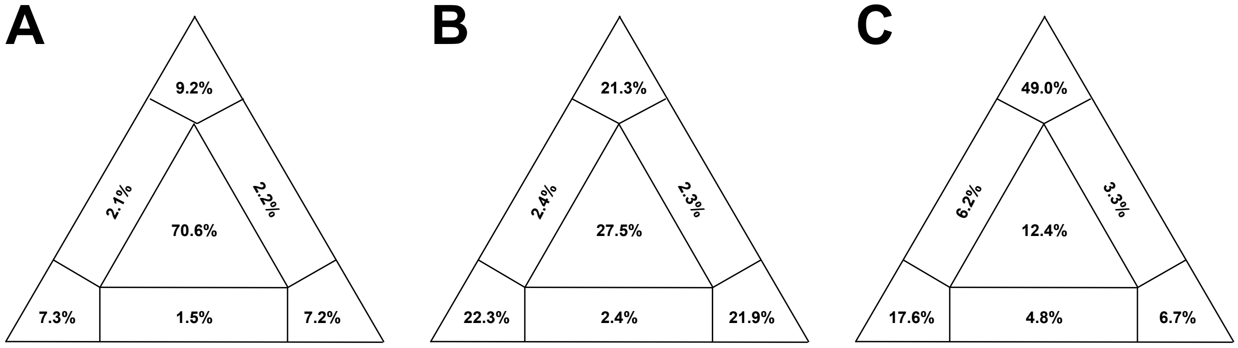


**Supplementary Figure 6. Likelihood mapping shows varying levels of information in outlying gene.** Quartet distributions of A) gene 7169_C12 from Wickett et al. (2014), B) gene 6349_C12 from Wickett et al. (2014) and C) gene 8314 from Chiari et al. (2012).
